# Improved diagnosis of viral encephalitis in adult and pediatric hematological patients using viral metagenomics

**DOI:** 10.1101/2020.06.05.136317

**Authors:** Ellen C. Carbo, Emilie P. Buddingh, Evita Karelioti, Igor Sidorov, Mariet C.W. Feltkamp, Peter A. von dem Borne, Jan J.G.M. Verschuuren, Aloys C.M. Kroes, Eric C.J. Claas, Jutte J.C. de Vries

## Abstract

Metagenomic sequencing is a powerful technique that enables detection of the full spectrum of pathogens present in any specimen in a single test. Hence, metagenomics is increasingly being applied for detection of viruses in clinical cases with suspected infections of unknown etiology and a large number of relevant potential causes. This is typically the case in patients presenting with encephalitis, in particular when immunity is impaired by underlying disorders.

In this study, viral metagenomics has been applied to a cohort of hematological patients with encephalitis of unknown origin.

Since viral loads in cerebrospinal fluid of patients with encephalitis are generally low, the technical performance of a metagenomic sequencing protocol enriched by capture probes targeting all known vertebrate viral sequences was studied. Subsequently, the optimized viral metagenomics protocol was applied to a cohort of hematological patients with encephalitis of unknown origin.

Viral enrichment by capture probes increased the viral sequence read count of metagenomics on cerebrospinal fluid samples 100 – 10.000 fold, compared to unenriched metagenomic sequencing.

In five out of 41 (12%) hematological patients with encephalitis, a virus was detected by viral metagenomics which had not been detected by current routine diagnostics. BK polyomavirus, hepatitis E virus, human herpes virus-6 and Epstein Barr virus were identified by this unbiased metagenomic approach.

This study demonstrated that hematological patients with encephalitis of unknown origin may benefit from early viral metagenomics testing as a single step approach.

**Highlights:**

- A metagenomics protocol employing virus capture probes was validated and retrospectively applied to 41 hematological adult and pediatric patients presenting with encephalitis of unknown aetiology
- Viral enrichment by capture probes increased sensitivity of viral metagenomics on cerebrospinal fluid samples 100 – 10.000 fold, compared to unenriched metagenomic sequencing
- In 12% of hematological patients with encephalitis of unknown origin, a virus was detected by viral metagenomics, which was not found by routine diagnostics
- Viral metagenomics represents a valuable addition to the diagnostics repertoire for hematological patients with suspected CNS infection

## Introduction

Encephalitis is an important clinical condition with high morbidity and mortality and therefore necessitates a proper and timely diagnosis and pathogen identification [1]. However, up to 63% of the encephalitis cases remain undiagnosed [1] and as a result, no targeted treatment can be initiated, no specific prognostic information can be obtained, and in outbreak settings no effective preventive measures can be taken.

Metagenomic next-generation sequencing has the potential to detect the full spectrum of viral pathogens in a single test. An increasing number of case reports have described the application of metagenomics to clinical cases of encephalitis of unknown origin in both immunocompetent and immunocompromised patients [2–15]. Immunocompromised patients are most at risk of infection with unexpected and novel pathogens and may present with insidious clinical symptoms [2, 16]. Recent prospective studies evaluated the use of viral metagenomics for undiagnosed cases in parallel with conventional diagnostics over a period of one year or longer [17, 18]. The minority of the patients included were immunocompromised, mainly due to HIV and solid organ transplants. To date, no metagenomic cohort studies have been published focusing on hematological patients with encephalitis.

Cerebrospinal fluid (CSF) remains the most common sample type obtained for diagnostics in cases of encephalitis, though brain biopsies tend to have a higher diagnostic yield of metagenomics [2, 10, 19, 20] as viral loads are lower in CSF. Moreover, metagenomic analysis is greatly affected by an extremely low pathogen-to-host genome ratio. Consequently, a lower sensitivity of metagenomic sequencing has been reported, when compared with conventional PCR-based molecular assays [21–26]. Host cell depletion is one way to increase the relative abundance of viral nucleic acids in metagenomic sequencing, but has not consistently been reported as beneficial when analyzing clinical samples [26]. In contrast, virus genome enrichment by means of capture probes has been shown to significantly enhance virus detection when sequencing for example respiratory samples [27–30].

In this study, the technical performance of a metagenomic sequencing protocol using capture probes targeting all known vertebrate viral sequences was determined when applied to CSF samples. This technical performance study was followed by a retrospective cohort study with hematologic adult and pediatric patients with encephalitis of unknown etiology.

## Methods

### Patient and sample selection

For the technical validation study, fifteen CSF samples of patients with encephalitis of known etiology previously sent to the Clinical Microbiological Laboratory (CML) of the Leiden University Medical Center (LUMC, The Netherlands) in the period of 2012-2017 were selected based on positive real-time PCR findings. These samples were tested by means of a lab-developed metagenomic protocol with and without viral capture probes. Additionally, three tissue biopsies from enteral origin were tested since brain biopsies were only limited available.

Following the technical validation, a cohort of 41 adult and pediatric hematological LUMC patients presenting with clinical symptoms of encephalitis was selected for retrospective analysis. Their CSF samples and brain tissue (one patient) were previously sent to the CML for routine diagnostics in the period of 2011-2019 and selected based on negative real-time PCR results for viral and bacterial pathogens.

### Ethical approval

This study was approved by the medical ethics review committee of the Leiden University Medical Center (CME number B19.021)

### Routine real-time PCR testing (PCR)

In the absence of relevant travel history, the laboratory-developed molecular real-time PCR panel for detection of pathogens in CSF consists of herpes simplex virus type 1 and 2 (HSV1/2), varicella zoster virus, enterovirus and parechovirus. In immunocompromised patients, the panel is expanded with Epstein Barr virus, human cytomegalovirus, JC virus and human herpesvirus type 6 (HHV-6), upon clinical request. These real-time PCRs are performed with internal controls for nucleic acid extraction and real-time PCR inhibition as published previously [31–37]. The initial diagnostic results were confirmed in this study by retesting (see table 1) to ensure the sample integrity after storage at – 80°C.

### Metagenomic next-generation sequencing (mNGS)

The metagenomics protocol used has previously been described and optimized for simultaneous detection of RNA and DNA targets [38, 39]. In short, internal controls, equine arteritis virus (EAV) for RNA and phocid herpesvirus-1 (PhHV) for DNA viruses were spiked into the clinical samples.

Subsequently, nucleic acids were extracted directly from 200 μl CSF sample using the MagNApure 96 DNA and Viral NA Small volume extraction kit on the MagNAPure 96 system (Roche Diagnostics, Almere, The Netherlands) with 100 μL output eluate. Extraction buffer only was used as negative control (for extraction, library preparation, and sequencing). From each sample 50 ul of eluate was used as input and concentrated using the SpeedVac vacuum concentrator (Eppendorf). Samples were dissolved in 10 μl of master mix for fragmentation (consisting of NEB next First Strand Synthesis, random primers and nuclease free water). RNA library preparation was performed using NEBNext Ultra II Directional RNA Library prep kit for Illumina with several in-house adaptations [39] to the manufacturers protocol in order to enable simultaneous detection of both DNA and RNA in a single tube per sample. Poly A mRNA capture isolation, rRNA depletion and DNase treatment steps were omitted, and diluted full size Y-shaped, dual indexed adaptors (1.5 uM) were used. For comparison, library preparation by means of the NEBNext Ultra II DNA Library preparation kit was performed with preceding cDNA and second strand synthesis step. Resulting amplified libraries were used as input material for capture of specific target regions or were subjected to sequence analysis without further processing.

Clustering and metagenomic sequencing using the NovaSeq6000 sequencing system (Illumina, San Diego, CA, USA) was performed according to manufacturer’s protocols. Primary data analysis and results Image analysis, base calling, and quality check was performed with the Illumina data analysis pipeline RTA3.4.4 and bcl2fastq v2.20. Approximately 10 million 150 bp paired-end reads were obtained per sample.

### Viral capture probe enrichment

The quality and quantity of the amplified libraries before capture were determined using the Fragment Analyzer (Agilent) and Qubit (Invitrogen) respectively. For capturing, 250 ng of four amplified DNA libraries were combined in a single pool resulting in a combined mass of 1 μg. For enrichment of the DNA sample library pools, the SeqCap EZ HyperCap Workflow User’s Guide (Roche) was followed with several in-house adaptations to the manufacturers protocol. Briefly, human Cot DNA and blocking oligos (Integrated DNA Technologies) were added to each library pool to block non-specific cross hybridization. The target regions were captured by hybridizing each pool of four sample libraries with the SeqCap EZ probe pool [40] overnight. The HyberCap Target Enrichment kit and Hyber Cap Bead kit were used for washing and recovery of the captured DNA. Finally, post-capture PCR amplification was performed using KAPA HiFi HotStart ReadyMix (2X) and Illumina NGS primers (5 μM), followed by DNA purification using AMPure XP beads. Quality and quantity of the post-capture multiplexed libraries were determined by Fragment Analyzer (Agilent) or Bioanalyzer (Agilent).

### Bioinformatic analysis

After quality pre-processing, sequencing reads were taxonomically classified with the pipeline Centrifuge [41] using an index database constructed from NCBI’s RefSeq and taxonomy databases (accessed April 4^th^, 2019). Reads with multiple best matches were uniquely assigned to the lowest common ancestor (k = 1 Centrifuge setting; previously validated [39]. Negative control sequence reads were subtracted from patient sample reads by Recentrifuge 0.28.7 [42]. Metagenomic findings were confirmed by a second pipeline, GenomeDetective [43] version 1.111 (accessed December 2018—January 2019) accounting for horizontal genome coverage (%) and confirmatory real-time PCR. Read counts were normalized for total read count and genome size.

## Results

### Technical performance on PCR-positive CSF samples

The results of the comparison of the metagenomic protocol with and without viral enrichment using capture probes for real-time PCR positive clinical CSF samples are shown in table 1. The metagenomic protocol without enrichment failed to detect the target viruses in three out of 18 cases. In contrast, the metagenomic protocol with enrichment for vertebrate viruses by capture probes detected all viruses that had been detected by real-time PCR. The target virus read counts were increased 100-10.000 fold after viral enrichment. Plots of horizontal coverage of viral sequences, with and without viral capture probes, are shown in Figure 1.

**Figure 1.**
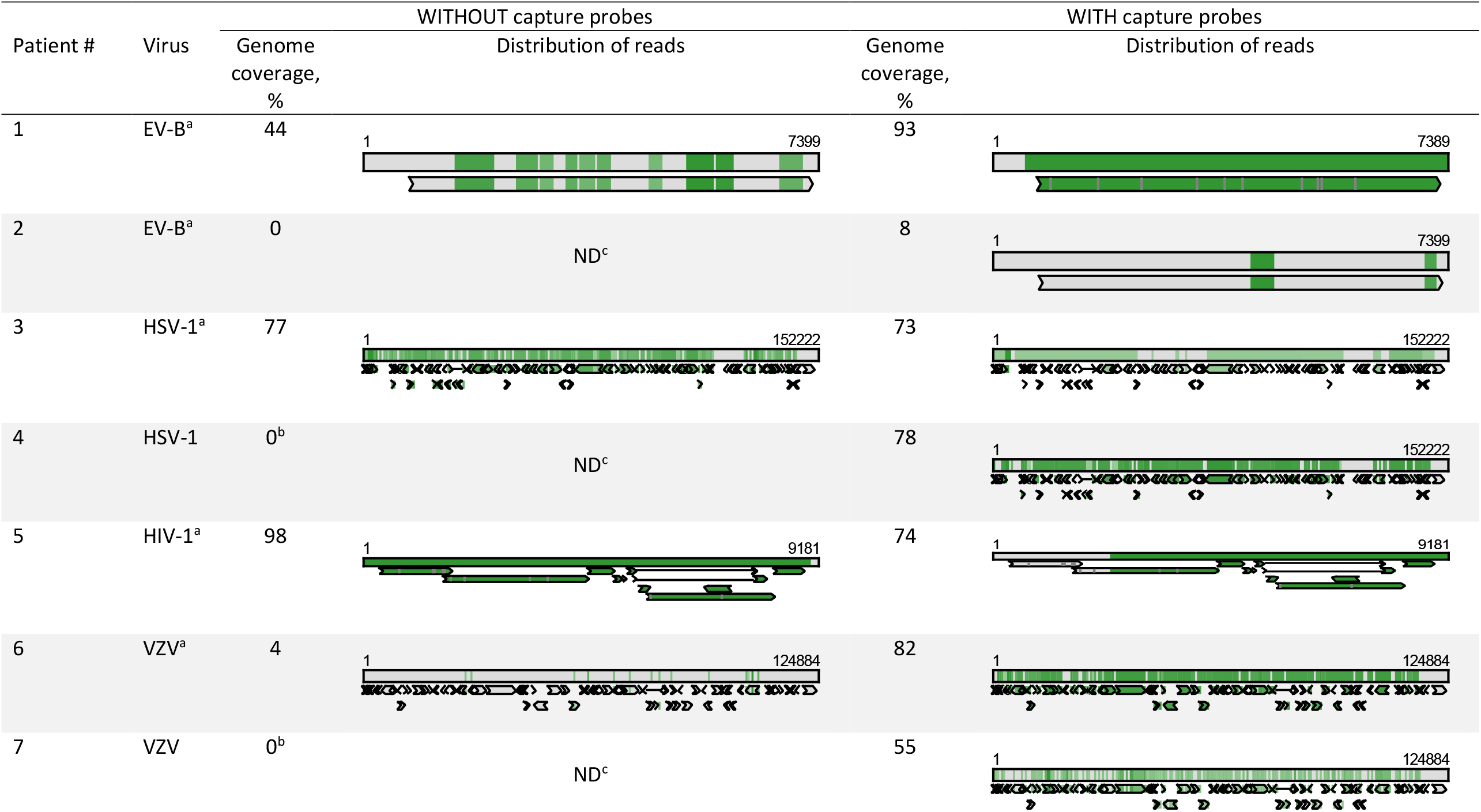
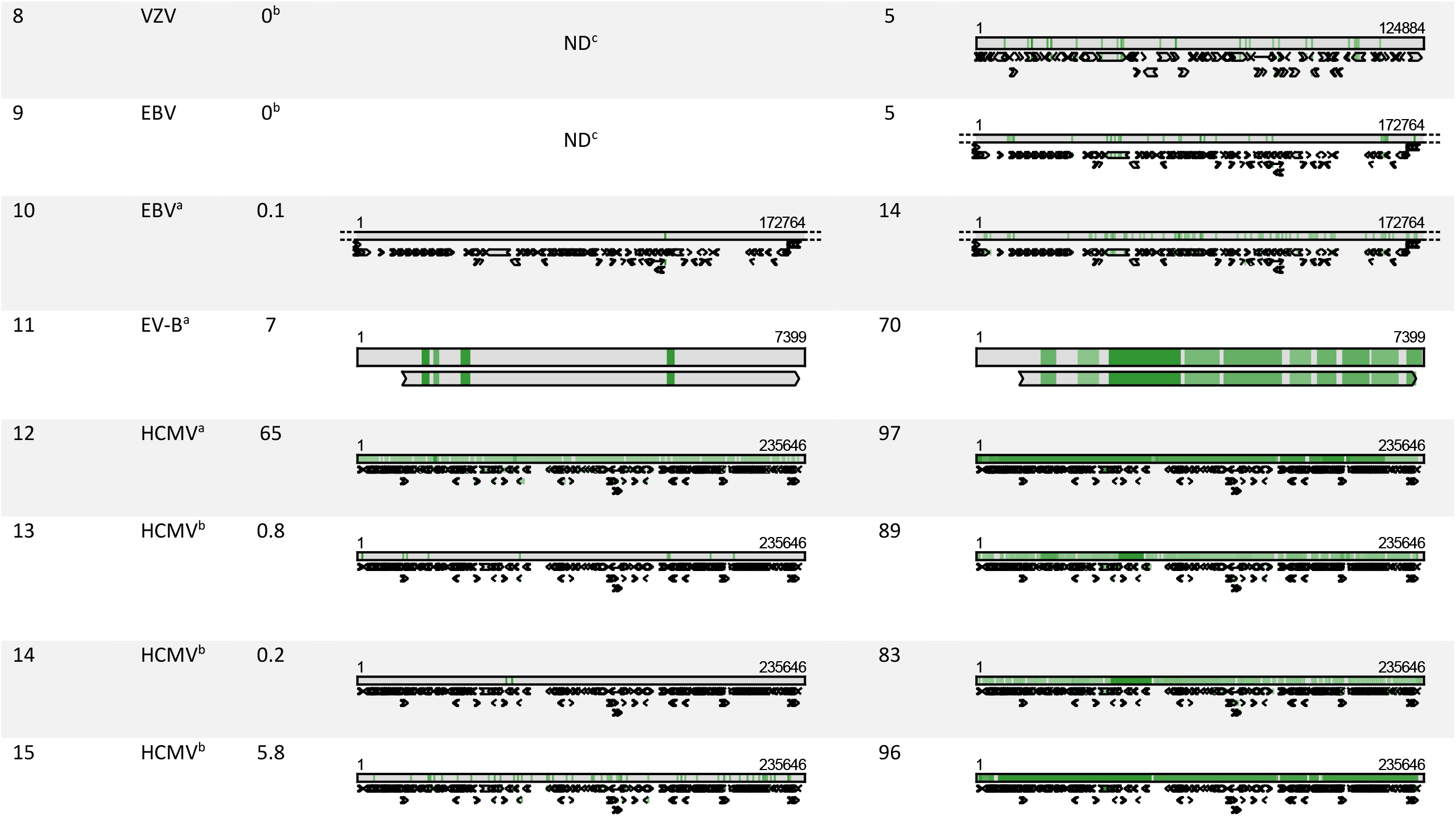
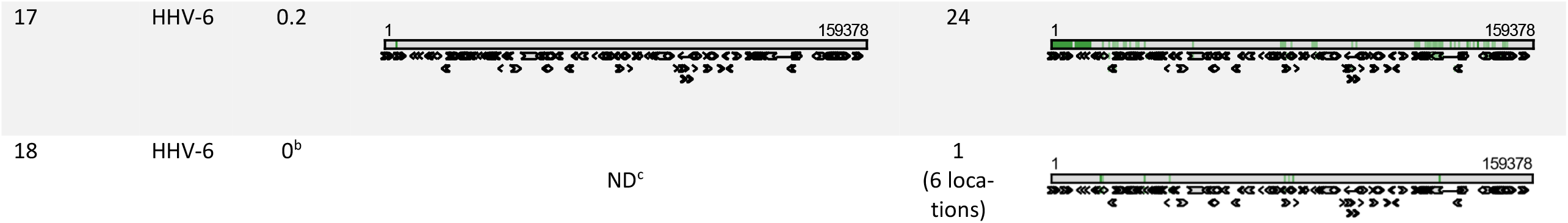
Horizontal genome coverage of PCR target viruses in technical performance study without (left) and with viral capture probes (right). Top bar represents nucleotide alignment, bottom bar(s) represents amino acid alignment, green zone: matching sequences. Sample 16 is not included because of negative PCR results. EV-B, enterovirus type B; HCMV, human cytomegalovirus; HSV, human simplex virus; VZV, varicellovirus; HIV, human immunodeficiency virus; ND, not detected ^z^ NEBNExt Ultra II Directional RNA Library preparation kit with in-house adaptations for total NA sequencing (see table 1 and methods) ^b^ NEBNExt Ultra II DNA Library preparation kit preceded by cDNA synthesis (see table 1 and methods) ^c^ Not detected (GenomeDetective)

**Table 1.** Comparison of the metagenomic protocol with and without viral capture probes in a panel of PCR positive CSF samples.

| Patient | Sample type | PCR result<br>(Initial Cq-value/load, retested value) | mNGS results,<br>without viral<br>probes (read count,<br>Centrifuge) |  | mNGS<br>results,<br>with viral<br>probes<br>(read<br>count,<br>Centrifuge)<br><sup>a</sup> | Increase<br>in read<br>count (-<br>fold) |
| --- | --- | --- | --- | --- | --- | --- |
|  |  |  | Adapted<br>RNA<br>prep.<sup>a</sup> | DNA<br>prep.<br>incl.<br>cDNA<sup>b</sup> |  |  |
| 1 | CSF | Enterovirus (27, 27) | 0 | 367 | 515.069 | 1.404 |
| 2 | CSF | Enterovirus (30, 34) | 0 | 0 | 12.368 | >12.368 |
| 3 | CSF | Herpes simplex virus type 1 (25) | 6.616 | 4.842 | 3.302.218 | 499 |
| 4 | CSF | Herpes simplex virus type 1 (30) | NT | 144 | 913.662 | 6.345 |
| 5 | CSF | HIV type 1 (302.500c/mL<sup>c</sup>) | 2.281 | 187 | 38.749.926 | 16.988 |
| 6 | CSF | Varicella zoster virus(27, 28) | 286 | 0 | 334.368 | 1.169 |
| 7 | CSF | Varicella zoster virus (30, 28) | NT | 36 | 131.138 | 3.643 |
| 8 | CSF | Varicella zoster virus (31, 31) | NT | 3 | 10.241 | 3.412 |
| 9 | CSF | Epstein-Barr virus (4.8, 4.3 log<sub>10</sub> IU/mL) | NT | 4 | 8.172 | 2.043 |
| 10 | CSF | Epstein-Barr virus (3.8, 4.1 log<sub>10</sub> IU/mL) | 0 | 90 | 28.044 | 312 |
| 11 | CSF | Enterovirus (33, 34) | 0 | 0 | 15.829 | >15.829 |
| 12 | Biopsy | Human cytomegalovirus (22, 23) | 2.228 | 8.000 | 2.047.002 | 256 |
| 13 | Biopsy | Human cytomegalovirus (22, 26) | NT | 193 | 169.154,<br>113.777<br>(duplicate) | 876 |
| 14 | Biopsy | Human cytomegalovirus (24, 28) | NT | 96 | 160.639 | 1.673 |
| 15 | CSF | Human cytomegalovirus, resistant (27) | NT | 22.350 | 3.577.617 | 160 |
| 16 | CSF | CSF: negative but biopsy astrovirus PCR positive | 0 | 0 | 0 | Not applicable |
| 17 | CSF | Human herpes virus type 6 (32, 26) | 26 | 306 | 168.837 | 552 |
| 18 | CSF | Human herpes virus type 6 (35, 34) | NT | 0 | 1.283 | >1.283 |

mNGS; metagenomic next-generation sequencing, CSF; cerebrospinal fluid, NT; not tested
<sup>a</sup> NEBNext Ultra II Directional RNA Library preparation kit with in-house adaptations for total NA sequencing (see methods)
<sup>b</sup> NEBNext Ultra II DNA Library preparation kit preceded by cDNA and 2<sup>nd</sup> strand synthesis for total NA sequencing (see methods)
<sup>c</sup> Insufficient material available for retesting

### Retrospective study: clinical cohort

Following the validation of the use of the viral capture probes, the metagenomic protocol was used for the clinical application study on samples of pediatric and adult hematological patients with encephalitis of unknown etiology. In total 46 samples (42 CSF samples, one brain biopsy, three blood samples) of 41 patients, including 17 children, were tested. Viral metagenomic sequencing resulted in virus detection in four CSF samples and one brain biopsy (5/41, 12%, Table 2). The clinical symptoms, underlying condition, imaging findings and treatment are shown in Table 3. In these five cases, the virus detected by means of metagenomics had not been targeted by the routine PCR assays that were performed initially.

**Table 2.** Findings by viral metagenomic sequencing of 41 pediatric and adult hematological patients in CSF and brain biopsy samples.

| Patient | Sample type | Initially requested molecular diagnostics (real-time PCR, Cq-value) | mNGS results (viral probe capture) | Read count (Centrifuge /Genome Detective)<sup>a</sup> | Genome coverage (Genome Detective) |  | Confirmatory testing (PCR) |
| --- | --- | --- | --- | --- | --- | --- | --- |
|  |  |  |  |  | % | Depth | Target PCR Cq/load, retested value/load |
| 1, child | Brain tissue, post-mortem | Brain biopsy: HSV-1/2, JC virus, enterovirus, parechovirus, HCMV, EBV, HHV6, mycobacterium tuberculosis, and Whipple's disease: negative.<br>Positive: adenovirus Cq 36, EBV Cq 37, HHV-6 Cq 33<br>CSF: Adenovirus, EBV, HHV-6, and VZV: negative | Biopsy: BK polyomavirus<br>CSF: negative | 140/<br>857 | 63 | 33 | BKPyV PCR positive<br>Cq 22/<br>4.031.000, 549.400 c/ml |
| 2, adult | CSF | HSV-1/2, VZV, enterovirus, parechovirus, HCMV, EBV, JC virus: negative | Human herpesvirus type 6<sup>a</sup> | 29.398/<br>225.466 | 32 (>30 regions) | 576 | HHV-6 PCR positive<br>Cq 26, 29 |
| 3, adult | CSF | HSV1/2, VZV, HCMV, toxoplasma: negative | Human herpesvirus type 6 | 1.117/<br>82.961 | 5 (>25 regions) | 1330 | HHV-6 PCR positive<br>Cq 30, 28 |
| 4, adult | CSF | HSV1/2, VZV, HCMV, EBV, adenovirus, HHV6, BK, JC, enterovirus, and parechovirus: negative | Hepatitis E virus<sup>a</sup> | 61/<br>2767 | 1 | 2690 | HEV PCR positive<br>Cq 36, 37 |
| 5, adult | CSF | HSV-1/2, VZV, HCMV, JC virus, adenovirus, enterovirus, parechovirus, m. pneumoniae, listeria, m. tuberculosis, and HHV-6: negative | Epstein-Barr virus<sup>a</sup> | 26618/<br>602.782 | 46 | 990 | EBV PCR positive<br>Cq 28/ 21.380c/ml |

Cq-value; quantification cycle value, mNGS; metagenomic next-generation sequencing, ped.; pediatric patient, HSV-1/2; herpes simplex virus type 1/2, HCMV; human cytomegalovirus, EBV; Epstein Barr virus, HHV-6; human herpes virus type 6, VZV; varicella zoster virus; BKPyV; BK polyomavirus, CSF; cerebrospinal fluid
<sup>a</sup> Initially not tested for by PCR but diagnosed with a delay of up to 2 weeks

**Table 3.**
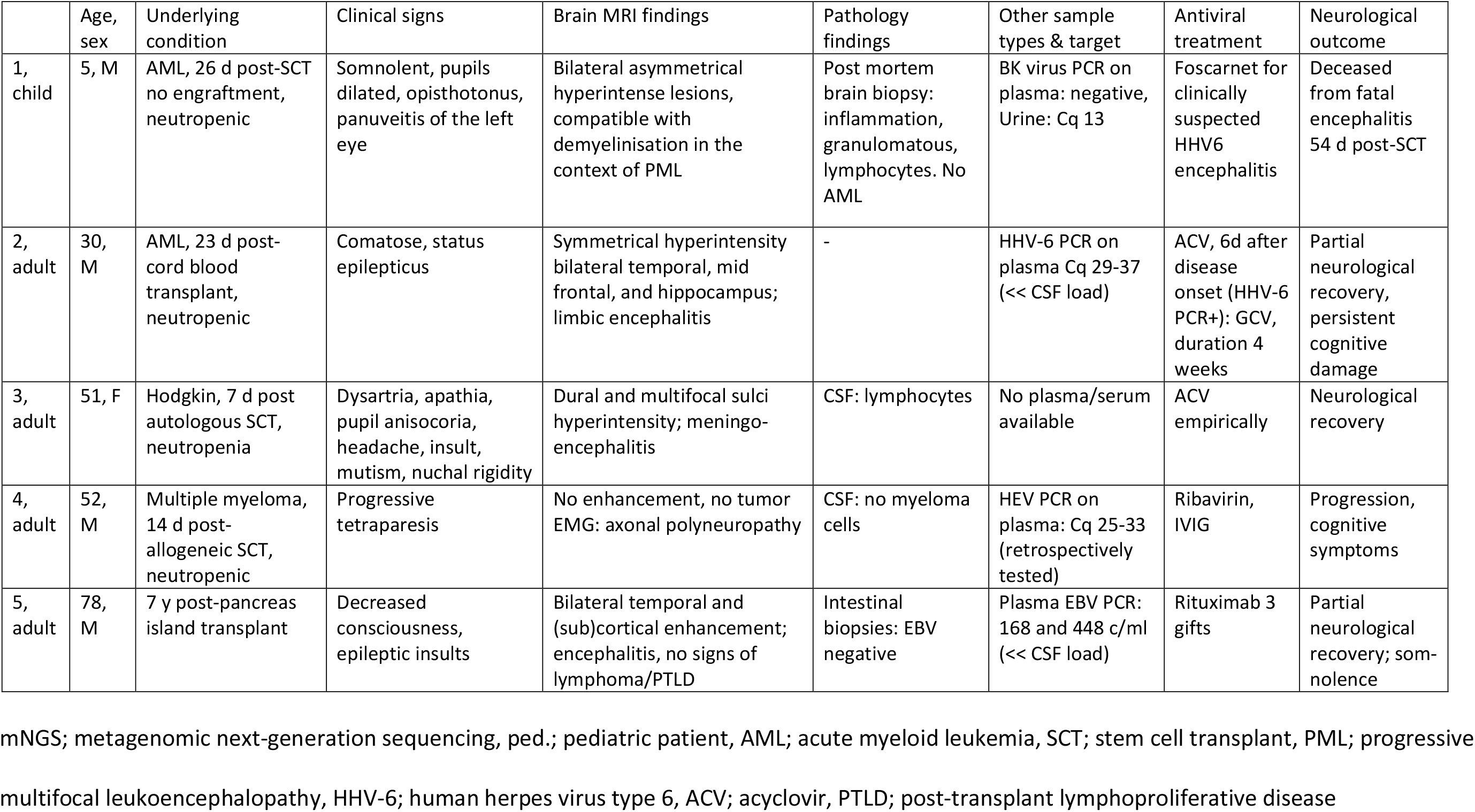
Clinical data of the patients with additional mNGS findings (see Table 2).

## Discussion

In this study, a metagenomic sequencing protocol employing virus capture probes was shown to be highly sensitive and of added value for detection of viruses when applied to a cohort of hematologic adult and pediatric patients with encephalitis of unknown origin. When compared to conventional molecular assays, viral metagenomic sequencing resulted in 12% additional findings, including some unexpected viruses initially not tested for.

An increase in the number of case reports involving the experimental use of metagenomic sequencing for diagnosing encephalitis in immunocompetent [3–6, 8, 11–13] and immunocompromised [7, 9, 10, 14, 15] patients is evident in recent literature [2, 44]. In these reported cases, the causes of encephalitis detected by metagenomic sequencing were novel, previously unknown viruses, but also, with similar frequency, well-established causes which could have been identified by conventional molecular techniques if only requested [2]. Other agents that were involved were known human pathogens that previously not had been observed as a causative agent of encephalitis [2]. Given the bias towards publication of cases with novel viruses, it is expected that when performing cohort studies, novel viruses will be less prominent, in line with our study. It must be noted that detection of novel viruses using a protocol employing virus capture probes is dependent on the amount of sequence similarity between novel and known viruses. None of the recent retrospective [45] and prospective [17, 18, 46, 47] cohort studies on metagenomic sequencing focused on neutropenic hematological patients, whom are likely at increased risk of infectious causes of encephalitis.

The clinical significance of detection of possibly latent and low level persistent viruses in CSF may be difficult to determine. Cohort studies do not provide the best support for causal relationships and the presence of the viruses detected in CSF in this study needs further investigation. For example, encephalitis caused by BK polyomavirus (BKPyV) has been indeed described in a series of case reports [48]. BK virus-associated progressive multifocal leukoencephalitis has previously only been reported in five cases [49]. In the current case, BKPyV was detected in brain tissue, which is considered the best support for diagnosing BKPyV virus encephalitis [49]. The absence of BK viremia in our case suggests localized reactivation of BKPyV in the central nervous sytem (CNS).

Likewise, positive findings of potentially latent viruses such as HHV-6 and EBV should be interpreted in the context of clinical presentation and sample type. HHV-6 DNA can be detected in the blood of approximately 50% of the hematopoietic stem cell transplant recipients [50], while the reported incidence of HHV-6 encephalitis is only 1% [51]. The presence of high viral loads in CSF when compared to blood, as seen in our cases of HHV-6 and EBV reactivation, is suggestive for localized CNS reactivation.

Hepatitis E virus (HEV) infection is associated with neurological dysfunctions, such as encephalitis and Guillain-Barré syndrome. This is supported by both clinical and laboratory studies, detecting HEV RNA in brain tissues of animals after experimental infection [52]. Neurological manifestations of hepatitis E virus infections are more frequently found in immunocompetent patients, suggesting pathophysiological mechanisms involving the immune response [53]. This may be the case in our patient given the lower viral load in CSF.

Though brain biopsies tend to have a higher diagnostic yield of metagenomics [2, 10, 19, 20], the most commonly acquired sample type in cases of encephalitis is CSF. Given the commonly low viral loads in CSF, optimal sensitivity is essential but challenging due to the high amount of background sequences [21–26]. Technical validation studies of viral metagenomic protocols using cerebrospinal fluid samples with known pathogens [25, 26, 54, 55] are essential to gain insight in its analytical performance including sensitivity. Virus enriched sequence analysis after probe capture has been shown to enhance virus detection significantly in respiratory samples [27–30, 56]. The current study confirms an increased sensitivity in CSF and tissue samples as well.

Summarized, the usefulness of viral metagenomics is dependent on several factors, including the technical aspects of the protocol, and the patient population studied. The current study shows that hematological patients may benefit from early, unbiased diagnostics by means of a virus enriched metagenomic sequencing protocol.

